# 3D Droplet-Based Bioprinting of Customized *In Vitro* Head and Neck Cancer Tumor Microenvironment Models

**DOI:** 10.64898/2026.03.27.714925

**Authors:** Victoria Messuri, Abbie Ha, Lissette A. Cruz, Daniel A. Harrington

## Abstract

*In vitro* models are increasingly critical for interrogating cancer biology and therapeutic response, however, accurately recapitulating the tumor microenvironment (TME) remains a persistent challenge, particularly in head and neck cancers (HNC) characterized by complex cell-matrix interactions and heterogeneity. Current models often lack independent tunability of biochemical and biophysical cues, limiting systematic investigation of microenvironmental cues in a high-throughput format.

Here, we establish a 3D droplet-based bioprinting platform for the fabrication of customizable *in vitro* TME models using poly(ethylene glycol) (PEG) hydrogels. Human HNC cell lines (FaDu and 2A3) with differing HPV statuses were bioprinted into PEG matrices spanning physiologically relevant stiffnesses (0.7–4.8 kPa) and compositions, including non-functionalized PEG and peptide-functionalized PEG (PEG^fnc^: RGD, YIGSR, CNYYSNS) and cultured for 7 days. Cluster growth, cell viability, and cluster morphology were assessed across multiple time points, matrix compositions, and matrix stiffnesses. Proliferation and endpoint phenotype expression were visualized using confocal microscopy through immunofluorescence.

Results indicated enhanced cell viability in PEG^fnc^ matrices, compared to non-functionalized matrices, while effect of matrix stiffness was less prominent. Median cluster size reached 40-50 μm by day 7, and linear mixed-effects modeling identified how changes in cluster surface area, volume, and tumoroid complexity varied with cell type, matrix, and stiffness. By decoupling and systematically varying key TME parameters, this approach provides a robust and scalable framework for dissecting tumor-matrix interactions and advancing physiologically relevant *in vitro* models for cancer research and therapeutic screening.

## 1. Introduction

With current recommendations from both the U.S. FDA and NIH in 2025 to shift drug screening research towards human-based non-animal models (NAMs), it is important to develop relevant, accurate, and replicable three-dimensional (3D) *in vitro* platforms for studying disease.^1,2^ Historically, models for studying cancer pathways and testing drugs have ranged from 2D monolayers on plastic to *in vivo* tumor studies in mice.^3^ Both extremes have strengths and flaws: 2D cell culture is inexpensive, simple, and fast, but cannot replicate complex multicellular organization; *in vivo* rodent studies offer complex physiologic replicas of native tissue, but are costly and not fully compatible with human cells. Over recent decades, the percentage of drugs advancing from successful preclinical studies to FDA approval has remained low due to poor translation into humans, which has been linked to the experimental models in which they were tested.^4–9^ Many high-throughput screens (HTS) for drugs rely on 2D cell cultures, which lack crucial cell-matrix interactions and the complex spatial arrangement of multiple cell types that are observed in 3D models.^10^ 3D “tumor engineered” hydrogel systems address the shortcomings of other models by enabling human cells to be encapsulated in miniaturized 3D mimics of the tumor microenvironment (TME), permitting cell migration, organization, and interaction with a surrounding extracellular matrix (ECM).^11^ Drug responses to hepatocellular carcinomas involving 6 different anti-cancer pharmaceuticals were shown to be significantly greater and wider in 3D aggregated spheroid models (containing a TME) when compared to 2D cultures, suggesting that 2D cultures and conventional spheroid systems are overly sensitive when measuring efficacy of drugs, and that more complex 3D systems (e.g., the incorporation of microenvironment components) improved the analytical process of discerning drug effectiveness.^10^ Several recent studies and reviews address continual improvements in 3D *in vitro* models for drug screening and their comparative advantages against 2D cultures.^12–16^ Synthetic hydrogels enable “bottom-up” modification of mechanical or biochemical characteristics through covalent control over crosslink density and integration of peptidic epitopes that influence cell adhesion or phenotype.

HTS for therapeutic compounds has been deployed successfully in 2D multi-well platforms, via developments in automated dispensing, liquid handling, and assays that enable fast readouts of simple viability indicators. High-content screening (HCS) combines the benefits of high-throughput drug screening (HTS) with cellular imaging for quantifiable data of complex systems.^13^ Although technology exists for spatial deposition of soft hydrogels within cells through 3D bioprinting, these elements have not yet merged in a significant way with HTS to enable complex experiments with discrete positioning of unique cell/gel combinations. Given that cancer biology involves more than just the cancer cell, a tailored approach that permits selective spatial deposition of multiple cell types and ECM components offers a better option for recreating key aspects of the TME.

In the present work, we apply these concepts to head and neck squamous cell carcinoma (HNSCC), the sixth most common cancer worldwide, representing nearly all cancers arising within epithelial linings of the oral cavity, throat, larynx, and nasal cavity. New diagnoses in 2022 reached approximately 760,000 worldwide with about 380,000 deaths.^17^ The causes of HNSCC are predominantly high tobacco and alcohol use, suspected germline mutations resulting in oral cavity and tongue tumors arising in younger patients, or human papillomavirus (HPV) infection.^18,19^ Collectively, HNSCC cases present the characteristics of late diagnosis, rapid progression, and resistance to treatments.^20^ After successful treatment by standard of care surgery and chemoradiation therapy, patients struggle with decreased quality of life, and there are notable failures of treatment especially in HPV-positive cases.^21^ Surgical procedures to remove tumors have been linked to immunosuppression, resulting in tumor recurrence.^21–25^ Although many 3D *in vitro* models have been developed for other cancer types, few have focused on HNSCC.^3,18^ Considering the prevalence and severity of this disease, it is crucial for high-fidelity tumor models to be designed and studied to answer biological questions and advance treatment options.

Here, we demonstrate human HNSCC 3D NAMs created by a droplet-based bioprinter (DB-BP) platform, which enables fast dispensing of cells suspended in clear hydrogels with spatial precision in multi-well formats designed for HTS/HCS (**Figure 1**). We explore the response of two immortalized human cell lines, reflecting HPV^+^ and HPV^−^ conditions, within hydrogels with variable gel modulus (0.7-4.8 kPa) and varied integration of peptidic ligands to promote cell adhesion or impact phenotype (RGD, YIGSR, CNYYSNS). These fibronectin, laminin β, and collagen IV-derived peptides support cell attachment, spreading, viability, and organization by providing key adhesive binding motifs and structural support. With a focus on these material properties, our quantified responses for cell viability, expansion, and tumoroid morphology in 3D, as a function of the above hydrogel variables, provide a roadmap for future implementation in drug screening applications.

**Figure 1.**
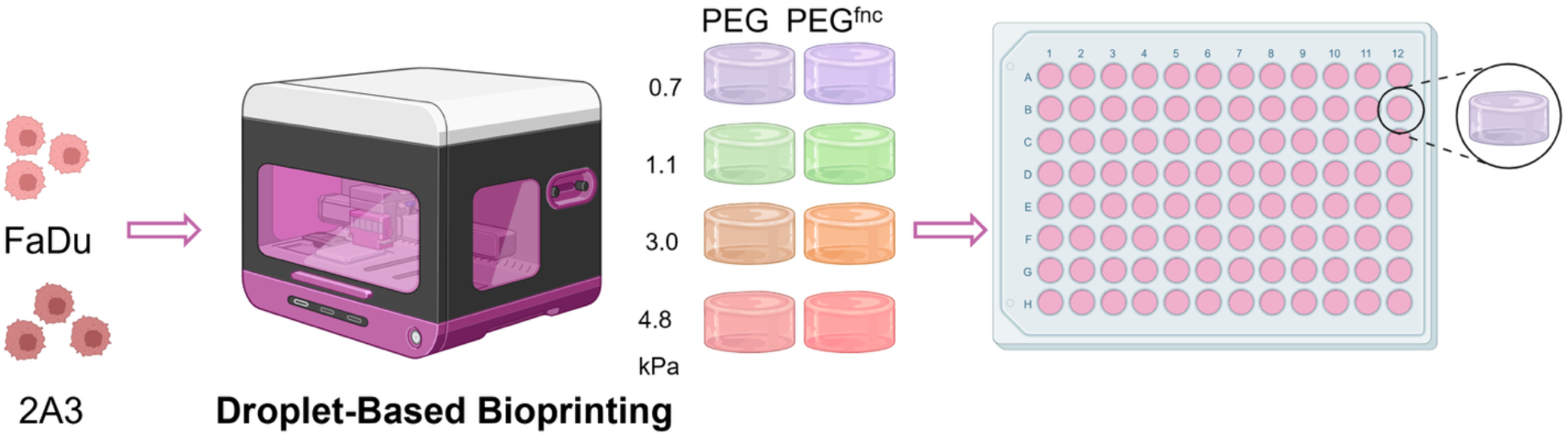
Printing Process used to Generate 3D Hydrogel Cancer Models. FaDu and 2A3 cells were suspended directly into a matrix metalloproteinase (MMP) labile bio-thiol activator which was then printed with its respective 4-arm maleimide PEG bioink to establish hydrogels upon component contact at room temperature. Both cell lines were printed into 8 different matrix conditions (PEG versus PEG^fnc^ in stiffnesses of 0.7, 1.1, 3.0, and 4.8 kPa) for a total of 16 matrix conditions investigated in this study. High-throughput printing was achieved through the use of 96-well plates, with the printing process allowing for spatial control of different cell and gel types to create a customized system based on user-preference.

## 2. Materials and Methods

### 2.1. Cell culture components and methods

FaDu (HTB-43) and 2A3 (CRL-3212) human-derived cell lines were purchased from ATCC, and cultured using ATCC-suggested media for each line. FaDu cells were originally derived from an Indian male with hypopharyngeal squamous cell carcinoma, and are HPV^−^. FaDu cells were cultured using ATCC-formulated Eagle’s Minimum Essential Medium (EMEM), Catalog No. 30-2003, with the addition of 10% (v/v) fetal bovine serum (FBS, R&D Systems, S111500) and 1% (v/v) penicillin-streptomycin (P/S). FBS was heat-inactivated (HI-FBS) prior to use by a standard treatment of 30 min at 56°C, followed by immediate quench in ice. 2A3 cells were derived from the FaDu cell line, but transduced to express HPV-16 E6/E7 genes, reflecting an HPV^+^ model.^26^ 2A3 cells were cultured in Corning Dulbecco’s Modification of Eagle’s Medium (DMEM) Reference No. 10-013-CV with the addition of 10% (v/v) HI-FBS, 1% (v/v) P/S and 0.2 mg/mL G-418. Trypsin/EDTA 0.25% (Corning 25-200-072) was used to passage cells by standard methods in 2D culture.

### 2.2. Droplet-Based Bioprinting procedures

A RASTRUM™ bioprinter (Inventia Life Sciences)^27–29^ was used for all bioprinting. Briefly, the RASTRUM deposits nanoliter-scale droplets, in a manner similar to that of an inkjet printer, onto standard multi-well cell culture plates within its own sterile benchtop cabinet. Hydrogels were generated in the instrument by automated sequential deposition of aqueous solutions of pH-adjusted 4-arm poly(ethylene glycol) (PEG) with terminal maleimide groups (“bioink”) and bifunctional bis-thiol PEG (“activator”) as ∼30 nL droplets onto the culture surface, where they crosslink immediately at room temperature via Michael addition to form 3D PEG-based hydrogel voxels,∼400 μm diameter.^30^ These voxels are deposited consecutively in multiple *xy* arrangements and stacked in *z*. To bioprint each cell type, the selected cell line was trypsinized from a tissue culture flask, neutralized in standard media, counted using a hemocytometer, pelleted at an appropriate concentration, and resuspended as a single-cell suspension in 200 μL of activator solution at 1 × 10^7^ cells/mL.

RASTRUM Cloud software was used to generate a PrintRun file and protocol for each cell line, hydrogel stiffness, and peptide composition. Prior to printing, the instrument was cleaned, primed, and calibrated for alignment following the manufacturer’s instructions. The printer was loaded with a sterilized flat, glass-bottom black-walled 96-well plate (Greiner Bio-One SensoPlate 655892) along with the RASTRUM cartridge that contained 40 mL sterile 18.2 MΩ•cm water, 24 mL 70% ethanol, 1.5 mL of “inert base” components, and 200 μL of the bioink and activators for the matrices. Two or three separate matrix conditions, at 20-24 replicates each, were printed per 96-well plate, and cell-laden hydrogels were printed in an “Imaging Model” configuration into the well plates, generating a centrally-positioned square hydrogel 500 μm high and 2.2mm in diameter. During printing, the instrument deposits the inert base layer of acellular hydrogel ∼220 μm thick, followed by the cellular layer, to ensure models are fully 3D. Cellularized hydrogel samples on a plate were typically printed within < 10 min. A Targeting Step was also completed during the print using a 96-well targeting plate to ensure accurate spatial deposition of the bioink and activator within the wells. Following completion of all prints on a plate, 200 μL of media was added to each well and replenished every two days. The printed specimens were incubated in standard HEPA-filtered humidified tissue culture incubators at 37°C/5% CO_2_/95% RH.

### 2.3 Hydrogel print parameters

Hydrogel precursor reagents varied according to the desired stiffness and composition. Each was identified as a two-part set of bioink and activator through RASTRUM Cloud, generously provided by Inventia Life Science. The matrices contained bioinks and activators (**Table 1**) to form both unfunctionalized PEG gels of stiffnesses 0.7, 1.1, 3.0, and 4.8 kPa (Px01.00, Px02.00, Px03.00, and Px06.00 respectively) and functionalized PEG gels containing the peptide sequences RGD, DYIGSR, and CNYYSNS (Px01.71, Px02.71, Px03.71, and Px06.71 respectively), each covalently bound to the matrix to produce concentrations of 1 mM in the final hydrogels, in the same four stiffness options. Reagents and basic chemistry were prepared as previously described.^28,29^ Activators contain an MMP-responsive bis-thiol that crosslinks with maleimide-functionalized PEG-based bioink; the Px06 (4.8 kPa) activator includes an additional 4-arm-PEG-thiol to increase stiffness. Prior published descriptions^28^ indicate 20 kDa PEG bioink, an MMP-cleavable peptidic activator (GCRDPLGLDRCG), and cysteine-terminal peptide epitopes.

**Table 1.** Compositions for Hydrogel Matrices.

| Stiffness (kPa) | Peptide Epitopes | Activator # | Bioink # | CAS # |
| --- | --- | --- | --- | --- |
| 0.7 | None | F176 | F38 | Px01.00 |
|  | RGD, DYIGSR, CNYYSNS | F176 | F320 | Px01.71 |
| 1.1 | None | F177 | F119 | Px02.00 |
|  | RGD, DYIGSR, CNYYSNS | F177 | F321 | Px02.71 |
| 3.0 | None | F178 | F97 | Px03.00 |
|  | RGD, DYIGSR, CNYYSNS | F178 | F322 | Px03.71 |
| 4.8 | None | F323 | F97 | Px06.00 |
|  | RGD, DYIGSR, CNYYSNS | F323 | F322 | Px06.71 |

All 16 experimental permutations were printed (**Table 2**), across the two cell types, four hydrogel stiffnesses, and two peptide compositions. 2-3 conditions were printed completely within a plate in a given day, using six batches to complete the series. Because samples did not repeat across batches, statistical analysis in R (lmer, below) was used to assess any confounding impact of batch on the measured outputs, and identified no statistically significant effect for log(area) or log(volume), but it did affect complexity, and was included in the model.

**Table 2.** Experimental Conditions.

| Cell Type | Stiffness (kPa) | Peptide Epitopes |
| --- | --- | --- |
| FaDu;<br>2A3 | 0.7,<br>1.1,<br>3.0,<br>4.8 | None;<br>RGD, DYIGSR,<br>CNYYSNS |

### 2.4. Cell viability and morphology assessment

Viability assays were performed by aspirating media from three wells from each condition, rinsing each well twice with Dulbecco’s phosphate buffered saline (DPBS, Cytiva SH30264.FS, with calcium and magnesium), and adding 200 μL of 2 μM calcein-AM (Biotium 80011-3), 2 μM ethidium homodimer-III (Biotium 40050) and 1 μg/mL Hoechst 33342 (Enzo Life Sciences ENZ-52401) in DPBS. Hydrogels at 1, 4, and 7 days after printing were incubated in the viability solution at 37°C for one hour then imaged on a Nikon C2 confocal microscope with 10x Plan Fluor objective, 1024×1024 resolution, in 200 μm Z-stacks with 5.2 μm step size. Light microscope images were captured daily on a Nikon Eclipse Ts2 microscope at 4x magnification.

Analysis of the Z-stacks for viability was performed using Bitplane Imaris v10. The Spots function was used with an estimated xy diameter of 10 μm and z diameter of 30 μm for all images. Spots were filtered by a quantity above an automatic threshold with background subtraction and counted. Percent viability was calculated based on the following equation:^31^

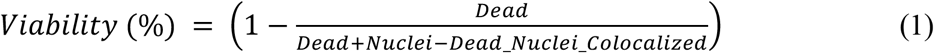

where “Dead” refers to the number of cells stained red by ethidium homodimer III, “Nuclei” refers to the number of cell nuclei stained by Hoechst, and “Dead_Nuclei_Colocalized” refers to cells that were stained both red and blue by ethidium homodimer III and Hoechst. Dual-stained cells were subtracted from the total spot count to avoid overcounting.

The viability Z-stacks were additionally used to determine measurements of surface area and volume for each cell cluster by using the “Surfaces” function in Imaris. Surfaces were created by creating imaginary surfaces engulfing the signal from the green (calcein-AM) channel, and assessed for the above parameters, excluding debris. The additional value of complexity was calculated as a description of the degree to which each cluster deviated from a perfect sphere, i.e. its production of irregular protrusions into the surrounding matrix, reflecting an invasive nature. Waddell’s traditional measure of sphericity^32^ is a common calculation for this, in which irregular non-spherical objects measure <1, and 1 reflects a perfect sphere. For statistical purposes, we defined complexity as an allometric residual:

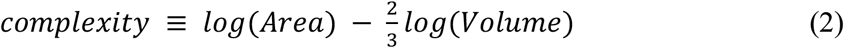

or effectively –log(sphericity). Complexity for a perfect sphere is ∼ 0.684, and all measures above that value reflect excess surface area beyond a sphere of the same volume.

### 2.5 Immunofluorescence assessment

Immunofluorescence (IF) staining for samples at day 7 of culture was performed by standard methods for 3D cultures, with details provided in Supporting Information. Each sample was aspirated, fixed, permeabilized, and blocked, with rinsing between steps. Samples were incubated overnight with primary antibodies (SI Table SI-1), rinsed with a wash buffer, incubated overnight with secondary antibodies and DAPI nuclear counterstain, rinsed, and stored in 0.1% sodium azide solution prior to imaging.

IF imaging was performed using a Nikon AX/AX R Confocal Microscope System, with NSPARC single-photon detectors and a 20x Plan Fluor objective. Images were captured with Resonant mode, Integration at 32×, using 405, 488, and 561 lasers, brightfield/transmitted light detector (TD), and standard filter blocks for DAPI, FITC, and TRITC. 3D z-stacks were 100 μm high, with 2.0 μm step size. 2×2 xy image stitching was utilized around the point with the highest concentration of in-focus cell clusters. Image analysis was performed using both Imaris v11 and Nikon NIS Elements.

### 2.6 Statistical Analysis

Statistical comparisons of viability for each matrix in Figure 3c were conducted by comparing means for each stiffness (PEG vs PEG^fnc^) via two-tailed, two-sample t-test assuming equal variances, with a significance level set at α = 0.05. Statistical significance is noted with *. Complementary comparisons of day 7 viability data across four stiffness levels at a single matrix composition were determined via one-way ANOVA followed by Tukey’s HSD post-hoc comparisons. All tests were two-tailed with the significance level set at α = 0.05. Statistical significance is noted by compact letter display above each bar. Graphs in Figure 3c reflect the mean of triplicate samples, and error bars reflect the 95% confidence interval (CI).

Measured values for area, volume, and complexity for each combination of cell type, matrix, and stiffness at day 7 were plotted as violin plots in GraphPad Prism 11. Each violin plot represented the entire set of clusters across three hydrogels. Cluster values for area and volume were log-transformed, fitted to a violin plot in Prism, then plotted on an antilog axis. Complexity values were fit directly in linear space and plotted on linear axes.

Cluster-level outcomes (area, volume, and a derived complexity metric) were analyzed using linear mixed-effects models in R with lme4 (lmer) to account for the hierarchical structure of clusters nested within gels/wells. For area and volume, responses were modeled on a centered natural-log scale (i.e., ln(area) and ln(volume)) to improve distributional behavior of positive, right-skewed size measures, address variance heterogeneity, and yield multiplicative interpretations on the original scale. Each model included a random intercept for well to capture within-well correlation among clusters and well-to-well heterogeneity. Batch-to-batch printing variation was evaluated as an additional random intercept. Batch did not improve fit for log(area) or log(volume) (Δχ^2^=0, p=1), but did improve fit for complexity (Δχ^2^=7.11, p=0.0077), so batch was retained for complexity only. Stiffness was treated as a continuous predictor on the centered log(kPa) scale. Fixed-effects were simplified using nested model comparisons fitted by maximum likelihood (ML) for valid likelihood-ratio tests (LRTs) when fixed effects differed, and final parameter estimates were obtained from the selected model refit using REML, consistent with standard mixed-model practice. Model form for the stiffness effect was selected per endpoint: we first tested linear vs quadratic (nested; ML fits; likelihood-ratio test), then compared quadratic vs natural-spline with df = 3 (non-nested; AIC/BIC), and finally confirmed day-7 adequacy by slice-level bias metrics (per-cell type × matrix), computed from fixed-effects predictions at the four stiffness points. This procedure supported a natural-spline stiffness effect (df = 3) for area and volume, and a quadratic stiffness effect for complexity.

To report biologically intuitive stiffness effects at the experimental levels (0.7, 1.1, 3.0, 4.8 kPa), we computed estimated marginal means (EMMs) and conducted Tukey-adjusted pairwise contrasts among these stiffness points within each cell type × matrix × time stratum using the emmeans framework (reference-grid predictions from the fitted model). Because the dataset was large, degrees-of-freedom adjustments were evaluated using the asymptotic option in emmeans to avoid the computational burden of Kenward-Roger/Satterthwaite methods at large N. For log-scale outcomes, contrasts were exponentiated to obtain ratios (and percent changes) on the original scale; under lognormal-like behavior, the back-transformed model mean on the log scale corresponds to the median on the original scale, supporting “typical” effect interpretation while enabling model-based inference.

## 3. Results

### 3.1. Cell growth and cluster count after bioprinting

Encapsulated cells in each hydrogel condition were imaged non-destructively by light microscopy over 7 days of culture (**Figure 2**). In addition, hydrogels were collected in triplicate at day 1, 4, and 7 of culture, stained with standard fluorescent viability stains Hoechst 33342, calcein-AM, and EthD-1 (identifying all cell nuclei, all live cells, and all compromised dead/dying cell nuclei, respectively), and imaged as z-stacks via confocal microscopy.

**Figure 2.**
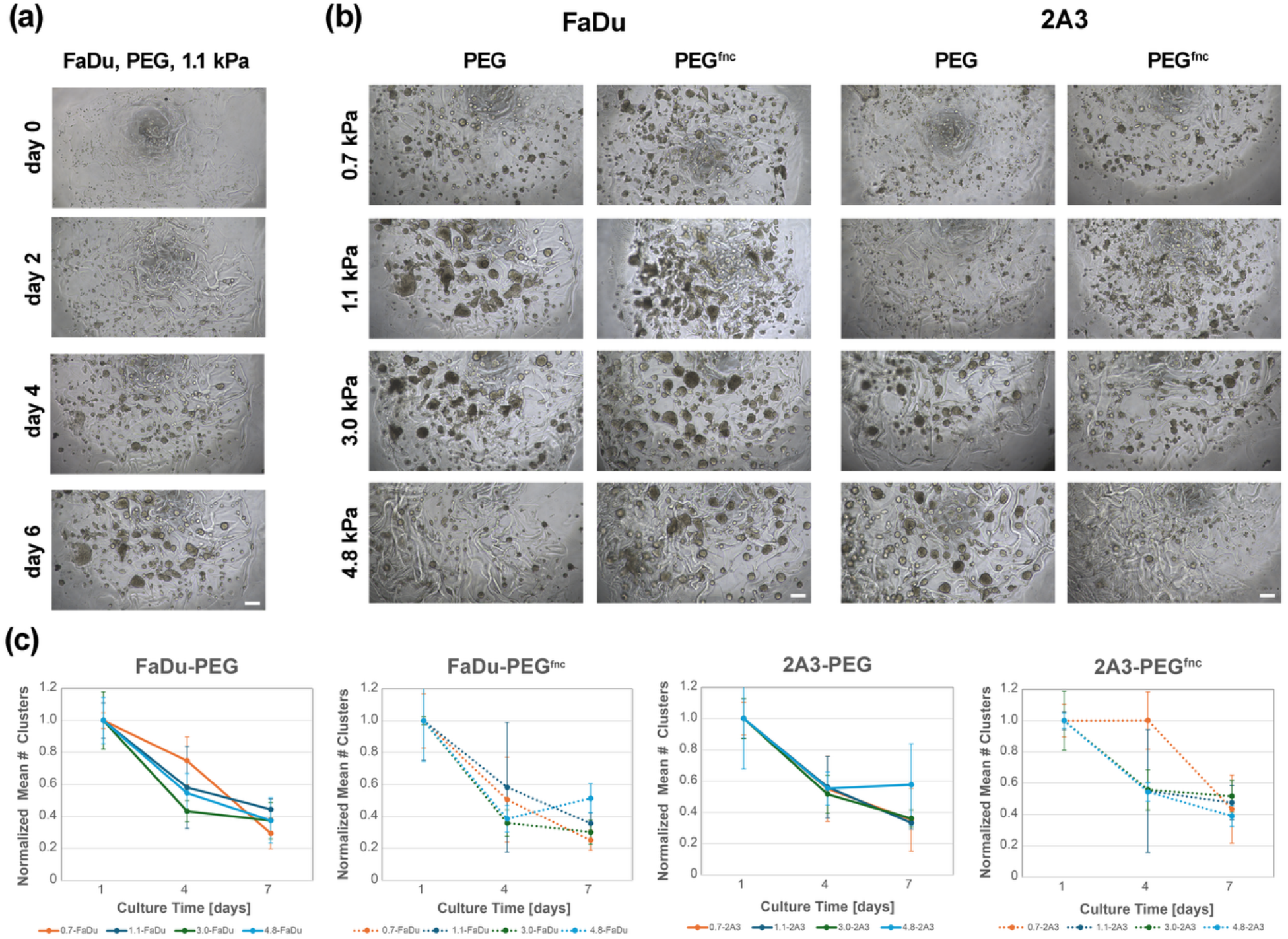
Hydrogels enable cell survival and rapid proliferation to form multicellular clusters. Light microscopy images of each cell type in each hydrogel condition were captured over 7 days to assess cell proliferation and morphology changes. (a) Images of FaDu cells, encapsulated as single cells in a PEG hydrogel of 1.1 kPa stiffness, at 0, 2, 4, and 6 days of culture show progressive cell proliferation and formation of multicellular clusters. (b) Final images of each cell type and gel condition, recorded at day 7 of culture, demonstrate the varied cell response to each condition. All samples were seeded initially as single-cell suspensions under identical cell concentrations. Corresponding day 0 images for panel (b) are provided in Supporting Information Figure SI-1. Scale bar = 100 μm. (c) Mean cluster counts for FaDu and 2A3 cells in each matrix, normalized to the day 1 count. Each point represents triplicate repeats, and error bars reflect the 95% confidence interval. Despite some variation, cluster counts are statistically indistinguishable for all conditions.

**Figure 3.**
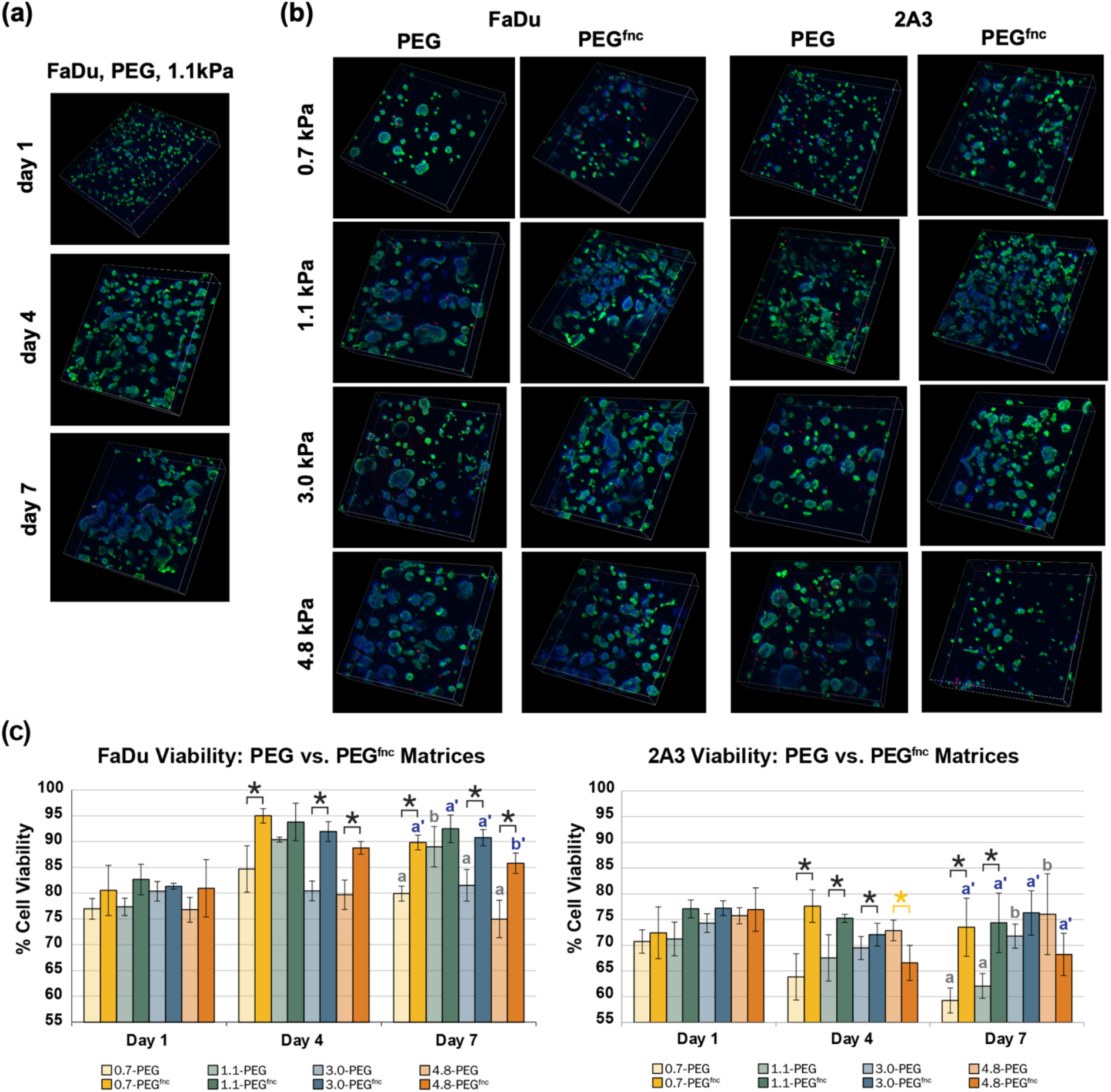
Viability stains demonstrate high viability of cell clusters over time. Hydrogel samples were collected at 1, 4, and 7 days of culture, stained to assess viability, and imaged via confocal microscopy as z-stacks. (a) Images of FaDu cells, in a PEG hydrogel of 1.1 kPa stiffness, at 1, 4, and 7 days of culture, show high cell viability (green) and few dead cells (red). The development of multicellular clusters over time tracks with observations from light microscopy in Figure 2. (b) Final images of each cell type and gel condition, recorded at day 7 of culture, reinforce the varied cell response to each condition. Scale: x and y dimensions reflect 826 × 826 μm, and z dimension reflects 200 μm depth. Blue: DAPI (nuclei), Green: calcein-AM (live cells), Red: ethidium homodimer (dead cell nuclei); (c) Quantified viability from reconstructed z-stacks for FaDu (left) and 2A3 (right) cells, printed in each hydrogel stiffness, either in PEG or peptide-functionalized PEG hydrogels. Each bar reflects triplicate specimens, error bars reflect 95% CI. Comparisons were conducted for cell viability between PEG and PEG^fnc^ matrices at each stiffness, * = p < 0.05. For Day 7 samples at the end of the experiment, letters identify statistically different results across four stiffnesses within the PEG (a, b) matrix or the PEG^fnc^ (a’, b’) matrix, α=0.05.

Each cell type was prepared as a single-cell suspension at an equivalent cell concentration. Day 0 images, recorded immediately after printing for each condition, are shown in Supporting Information **Figure SI-1** and confirm an essentially single-cell condition across the samples. Rapid cell proliferation to form multicellular structures was observed under every condition. **Figure 2a** demonstrates example growth for the FaDu cell line, encapsulated in PEG hydrogel, 1.1 kPa stiffness, at 0, 2, 4, and 6 days of culture. Multicellular clusters were observed within 24-48 hours of printing and occurred through a combination of early cell-cell adhesion and concurrent proliferation. **Figure 2b** shows the 16 conditions after 7 days of culture, with a range of cluster morphologies and sizes. By 7 days, most of the FaDu multicellular cultures appeared qualitatively larger in size than the 2A3 clusters in most conditions.

Through imaging of the fluorescent clusters at days 1, 4, and 7, and digital quantification of cluster number in Imaris, the relative change in the mean number of clusters in each matrix could be determined. As shown in **Figure 2c**, the number of clusters, normalized to the initial value at day 1, decreased over time for all matrix conditions and cell types, following a trend that was independent of cell type, matrix stiffness, or matrix composition.

### 3.2. Cluster viability after bioprinting

Confocal imaging of the growing cell clusters, stained with fluorescent viability markers, confirmed observations from light microscopy. Rapid cluster growth over 7 days was observed (example for FaDu cells in 1.1 kPa PEG hydrogels shown in **Figure 3a**), and images at day 1 suggested that the printing process did not yield significant distress onto the cells. Images on day 7 (**Figure 3b)** demonstrate the cells’ progressive growth into multicellular clusters. Digital quantification of these image stacks identified that the FaDu cell line retained higher viability in the hydrogels as compared to the 2A3 cell line (**Figure 3c**). Both cell lines exhibited ∼75% viability on day 1, but the FaDu cell line showed a trend for increasing viability for most peptide-functionalized matrix conditions over time by days 4 and 7, while the 2A3 cell line showed an opposite trend of slightly decreasing viability across most conditions by days 4 and 7. In most conditions, the peptide-functionalized matrices supported higher cell viability at days 4 and 7, when compared to the same conditions in the unfunctionalized matrix.

Statistical analyses of all data sets confirmed that the most significant differences were observed at day 7, so statistical comparisons of stiffness effects focused on this timepoint. ANOVA comparisons of viability for the day 7 specimens within a given matrix composition, but across stiffnesses, were conducted (**Figure 3c)**. In general, viability was only modestly affected by matrix stiffness, except for the condition of 2A3 cells encapsulated within PEG hydrogels, which showed a trend of increasing viability with increasing matrix stiffness.

### 3.3. Cluster morphology measures

Confocal z-stacks for viability were reconstructed in 3D within Imaris, using the fluorescent signal from calcein-AM (green) to identify cell clusters, and each cluster was wrapped with a virtual surface using the Surfaces feature. Surface area and volume outputs were recorded for every cluster within a hydrogel, across all culture times and hydrogel conditions, yielding n∼200-500 clusters per specimen at day 1, and decreasing to n∼50-200 clusters per specimen by day 7, following the trends in **Figure 2a**. These area and volume measures additionally were used to calculate complexity (**eqn (2)**), a log-inverse of sphericity, as a proxy for the invasiveness of a cluster into the surrounding matrix. Nearly all samples had n=3 repeat specimens, except as noted.

**Figure 4 (a-c)** shows the measured median cluster surface area, volume, and the calculated complexity, respectively, for each combination of cell type, matrix, and stiffness level after 7 days of culture. Qualitative review of these data suggested that the replicates were largely reproducible within a condition. Although cluster data was preserved within a replicate for modeling purposes, we loosened this restriction to obtain a broader view of cluster populations as shown in the violin plots in **Figure SI-2**. These violin plots again identify the median of each population at day 7, with upper and lower quartiles. Because of the substantial spread in area and volume data across multiple decades, these data were log-transformed first, fit with the kernel density estimation to generate the violin outline, then replotted on an antilog scale to avoid compression. The medians and trends from these full data sets mirror the replicate-isolated results in **Figure 4**. Cutoffs near 100 μm^2^ for surface area were set to eliminate debris below single-cell size. These plots identify a median cluster size of ∼40-50 μm diameter after 7 days of culture, with a progressive loss of small/single-cell structures, with the exception of a bimodal response for the 2A3-PEG-4.8 condition, and a possibly similar response for the FaDu-PEG^fnc^-4.8 system. Complexity indices were higher for FaDu-containing samples, but all populations retained a significant proportion of nearly-spherical multicellular clusters.

**Figure 4.**
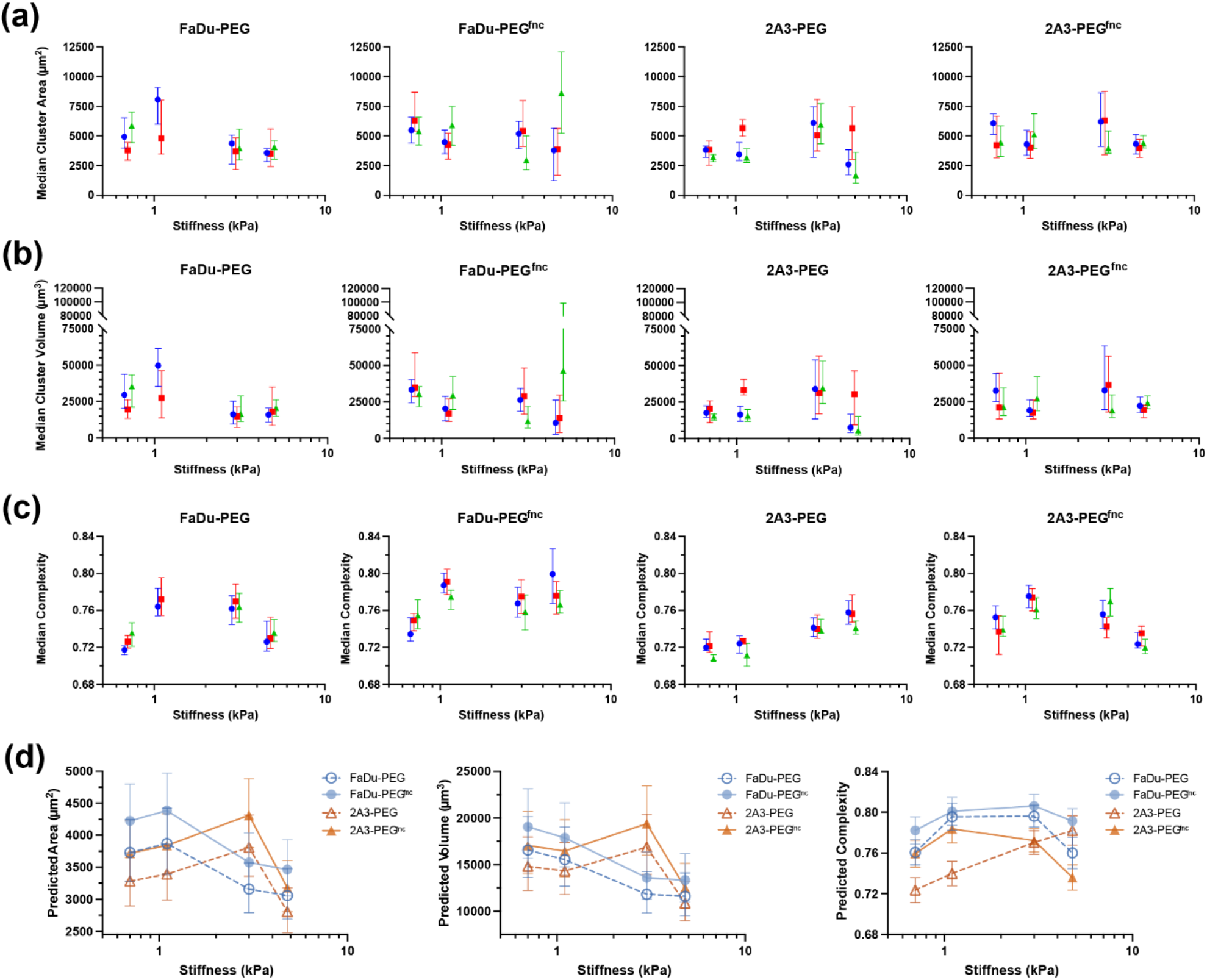
Cluster morphology measures and modeling at day 7 timepoint identify impacts of cell type and matrix parameters. 3D reconstructions of calcein-AM green-stained cell clusters for each cell type, matrix stiffness, and composition permutation were analyzed in Imaris to obtain per-cluster measurements of (a) surface area and (b) volume, and generate (c) complexity as a measure of invasiveness. (a-c) Graphs show the median of each value, ± 95% CI, n= 2-3 replicate gels (red, blue, green, offset for visibility) per condition, and n∼50-200 clusters per gel replicate. (d) Population data were modeled using lmer to generate plots of predicted cluster area, volume, and complexity as a function of stiffness. Cluster area and volume measures at day 7 varied with both stiffness and cell type and were cross-correlated. Area and volume magnitudes were correlated with matrix composition at day 7, but their change with stiffness was independent of matrix. Predicted complexity values aligned for FaDu cells in both matrices and for 2A3 cells in PEG^fnc^, but with unique behavior for 2A3 cells in PEG.

In an effort to quantify these observations and identify the contributions from each parameter, we used linear mixed-effects models (lmer) in R software. We modeled the raw data to identify the impact of fixed effects (cell type, matrix type, matrix stiffness, culture time) on cluster outputs, while controlling for random effects, such as intraspecimen variance and batch effects. Because this population data would be progressively right-skewed as clusters grow in the hydrogel, we elected to (a) model and report the median (as in **Figure 4, SI-2**) for each variable output, as the median is less prone to overinfluence by a few large outliers than the mean, and (b) model log transforms of area and volume, to stabilize variance of each dataset, closer to a normal distribution.

## 4. Discussion

As molecular modeling and drug design continue to improve at a rapid pace, there remains a need to test drugs in clinically relevant models, and with high efficiency. Deposition of cells within a 3D matrix, compatible with HTS/HCS assays, is a developing field, and the customization of methods to take advantage of that 3D environment is still in its early stages. For HNSCC, only a modest body of work has been published to extend examples for representative testing of human-derived cells in an efficient format.^3,18,33–35^ In the present work, we sought to make an initial contribution to this work through demonstration of two foundational cell lines, FaDu and 2A3 cells, in a novel DB-BP bioprinting system. The PEG-based hydrogel system has seen ample application for a wide range of 3D cell culture targets, particularly with the inclusion of biorelevant peptide epitopes and MMP-degradable crosslinkers. Recent publications in the context of the present bioprinting system support its potential for modeling a range of cell types in similar matrices.^36,37^

In our study, we found that both FaDu and 2A3 could be printed reliably in several matrix combinations. Even when printed in single cell suspensions, cells quickly proliferated in the 3D matrix and developed into multicellular clusters with median size ∼35-40 μm diameter, correlating empirically as clusters of ∼25-35 tightly-packed cells in 3-4 layers. Of course, a significant proportion of the cluster populations were far larger than this, and clusters with this level of mature development will have sufficient structure to retain diffusion gradients that impact drug diffusion in a clinically relevant way.

Despite notable changes in hydrogel stiffness and epitope content, we found that trends in the normalized mean number of clusters aligned onto a consistent behavior for both cell lines, across nearly all conditions (**Figure 2c**). Other literature studies have described excellent methods for assessing spheroid geometries within hydrogels,^38^ or have tracked cancer cell progression in alternate hydrogels at the per-cell level or full population level (e.g. via MTT assay).^39^ but we have not seen similar universal responses for the change in mean number of clusters. For drug screening use, this has relevance in retaining a sufficient number of viewable objects for statistical significance.

Multiple authors have described the potential impact of maleimide-thiol reaction kinetics on bulk hydrogel crosslink heterogeneity and encapsulated cell viability,^40–42^ however this speed was an enabling technology for the drop-wise bioprinting approach used in our work. Initial cell viability on day 1 after printing ranged from 71 to 83%, providing insight that the printing process itself allows for acceptable viability, which increases as cells are cultured within the hydrogels and are able to recover nutrients by the addition of media and proper incubation conditions. Although 2A3 cells demonstrated slightly lower average viability than FaDu cells on day 1, the two groups were largely within statistical error. However, a deviation between the two over time was observed, as FaDu viability increased over 4 and 7 days of culture, while 2A3 viability generally decreased. An important caveat to this is that the peptide-functionalized PEG^fnc^ matrices almost always supported significantly higher viability for both cell types at later timepoints when compared to PEG matrices of the same stiffness. Notably, 2A3 viability at 7 days was highest in the stiffest 4.8 kPa matrix; this correlated with atypical shift to the highest complexity. This result may suggest that cancer cells utilize high matrix stiffnesses for their benefit of remaining viable and/or invasive, as has been theorized for many other organ systems. The impact of the HPV type 16 E6 and E7 genes in the 2A3 line remains difficult to determine, and the above results add to the existing conclusions that the presence of HPV within HNSCC leads to increased difficulty in modeling.^18^

Further quantification of the viability images revealed trends in cluster formation and size. Broadly, the trend of cluster number reduction and condensation was remarkably consistent across the cultures. Thirteen out of the 16 conditions showed a decrease in the number of singular objects within the z-stacks throughout the culture period. Imaris 10 software was used to count individual objects using the Surfaces function, and data for each condition was normalized to its respective day 1. In the FaDu cells, the number of objects increased from day 4 to day 7 in the 4.8 kPa tripeptide condition, while the rest of the FaDu conditions displayed a decrease across the culture period. The 4.8 kPa blank condition in the 2A3 cells additionally showed an increase from day 4 to 7, though this finding was not statistically significant. Further, the 0.7 kPa tripeptide 2A3 condition showed no difference in number of objects from days 1 to 4, which is consistent with the light microscope images taken of the 0.7 kPa 2A3 conditions showing less and smaller cluster formation than those of FaDu and higher stiffnesses within both conditions.

Common questions for *in vitro* cancer models are “how big do the clusters become? how fast do they grow?” Answering these questions required treating the samples as populations of clusters, each with its own area/volume/complexity output. Modeling this cluster growth, and the details of cluster morphology, required using a lmer model, and adjusting the data with a log transform for area and volume to account for the increasingly right-skewed datasets as a tiny subset of clusters accelerated faster than the others. We additionally considered using Wadell’s sphericity as an output; its metric varies from 0 to 1, in which 1 represents a perfect sphere, and decreasing values reflect increasingly irregular objects. However, these data were again skewed, with an upper boundary and tail asymmetry, so we instead used complexity – the inverse log of sphericity – to generate a dataset that had no upper boundary, and was inherently log-transformed, more amenable to Gaussian modeling. In plain terms, higher complexity values identify “excess surface area” for an object of a given radius, reflecting irregularity and surface convolution. In our interpretation, this was a preferred viewpoint, rather than the “sphere-like compactness” measure of sphericity. The resulting models in **Figure 4c**, and the terms that were adjusted in **Table SI-2** to build them enabled us to identify trends and statistical interactions. For example, the consistent effect of functionalized matrix alone in developing a larger area and volume was notable, but this effect was only minimally cross-correlated with matrix stiffness in affecting both metrics. The value of this method was further revealed in the measure of complexity, for which 2A3 clusters in PEG matrices behaved substantially differently than the other three cell-matrix permutations, requiring a full 4-way model.

Synthetic polymers have been reported to comprise only 14% of papers between 2004 and 2020 to investigate matrix interactions with cancer.^43^ However, the utilization of synthetic polymers, such as PEG, allows for the study of stiffness with *in vitro* matrix systems *independently* due to the lack of receptor-binding sequences that are present in natural polymers^44^ yielding them better systems to investigate mechanical strength and offering more controlled degradation rates.^3^ The use of this synthetic polymer allowed for the incorporation of MMP-degradable peptide crosslinkers which contributed physiological relevance while also allowing for the unique control over other matrix parameters such as functionalization and stiffness.

A key advantage to droplet-based bioprinting is the ability to spatially sequester the multiple unique cell types that comprise the tumor microenvironment, within a HTS format. Prior work from other researchers has demonstrated the value of this strategy, culturing HNSCC lines like FaDu in 3D environments with associated fibroblasts.^45^ In future efforts, we look forward to building on the foundation set in the present work, leveraging this multicellular capability alongside our observations for matrix properties.

## 5. Conclusions

The method of droplet-based bioprinting to encapsulate live cells proved to be a feasible platform for the development of 3D *in vitro* systems for the application of disease study. Cancer clusters after 7 days of culture were viable, proliferative, and impacted by matrix parameters. Matrix customization with ECM-derived peptides and adjustable stiffness allows for a more physiologically relevant system when compared to 2D studies, with future opportunities for co-culture with cancer-associated fibroblasts and immune cells. This platform opens the potential for direct patient biopsies of HNC to be printed and analyzed for optimal personalized treatment conditions.

## Acknowledgements

This work was funded in part by a grant from VICTORY Houston. A.H. recognizes support from the Rice University Office of Undergraduate Inquiry. Confocal microscopy and digital quantification were performed using instruments and resources at the Center for Craniofacial Research Instrumentation Core, School of Dentistry, UTHealth Houston. The authors thank Julian (Nat) Holland for statistical input, and Inventia Life Science for use and support of the RASTRUM bioprinter, particularly Thomas Grundy, Whitney Symons, and Dwayne Dexter. We also thank Danielle Wu, Mary (Cindy) Farach-Carson, Daniel Carson, Caitlynn Barrows, Simon Young, Neeraja Dharmaraj, Max DeLeon, and Claudia Villalobos Perez for scientific input and feedback.

## Notes

### Competing Interest Statement

The RASTRUM instrument and associated consumables were provided to the laboratory as part of a trial agreement by Inventia Lifesciences. Although Inventia provided technical consultation on their workflows as needed, they had no scientific influence on the outcome of this work.

## References

(1) Commissioner, O. of the. FDA Announces Plan to Phase Out Animal Testing Requirement for Monoclonal Antibodies and Other Drugs. FDA. https://www.fda.gov/news-events/press-announcements/fda-announces-plan-phase-out-animal-testing-requirement-monoclonal-antibodies-and-other-drugs (accessed 2025–08-29).

(2) NIH to prioritize human-based research technologies | National Institutes of Health (NIH). https://www.nih.gov/news-events/news-releases/nih-prioritize-human-based-research-technologies (accessed 2025-08-29).

(3) Bonartsev, A. P.; Lei, B.; Kholina, M. S.; Menshikh, K. A.; Svyatoslavov, D. S.; Samoylova, S. I.; Sinelnikov, M. Y.; Voinova, V. V.; Shaitan, K. V.; Kirpichnikov, M. P.; Reshetov, I. V. Models of Head and Neck Squamous Cell Carcinoma Using Bioengineering Approaches. Crit. Rev. Oncol. Hematol. 2022, 175, 103724. 10.1016/j.critrevonc.2022.103724.

(4) Kola, I.; Landis, J. Can the Pharmaceutical Industry Reduce Attrition Rates? Nat. Rev. Drug Discov. 2004, 3 (8), 711–716. 10.1038/nrd1470.

(5) Hutchinson, L.; Kirk, R. High Drug Attrition Rates—Where Are We Going Wrong? Nat. Rev. Clin. Oncol. 2011, 8 (4), 189–190. 10.1038/nrclinonc.2011.34.

(6) Van Norman, G. A. Limitations of Animal Studies for Predicting Toxicity in Clinical Trials. JACC Basic Transl. Sci. 2019, 4 (7), 845–854. 10.1016/j.jacbts.2019.10.008.

(7) Errington, T. M.; Mathur, M.; Soderberg, C. K.; Denis, A.; Perfito, N.; Iorns, E.; Nosek, B. Investigating the Replicability of Preclinical Cancer Biology. eLife 2021, 10, e71601. 10.7554/eLife.71601.

(8) Marshall, L. J.; Bailey, J.; Cassotta, M.; Herrmann, K.; Pistollato, F. Poor Translatability of Biomedical Research Using Animals — A Narrative Review. Altern. Lab. Anim. 2023, 51 (2), 102–135. 10.1177/02611929231157756.

(9) Frühwein, H.; Paul, N. W. “Lost in Translation?” Animal Research in the Era of Precision Medicine. J. Transl. Med. 2025, 23 (1), 152. 10.1186/s12967-025-06084-3.

(10) Lee, S.-Y.; Hwang, H. J.; Lee, D. W. Optimization of 3D-Aggregated Spheroid Model (3D-ASM) for Selecting High Efficacy Drugs. Sci. Rep. 2022, 12, 18937. 10.1038/s41598-022-23474-5.

(11) Foglizzo, V.; Cocco, E.; Marchiò, S.; Foglizzo, V.; Cocco, E.; Marchiò, S. Advanced Cellular Models for Preclinical Drug Testing: From 2D Cultures to Organ-on-a-Chip Technology. Cancers 2022, 14 (15). 10.3390/cancers14153692.

(12) Mittler, F.; Obeïd, P.; Rulina, A. V.; Haguet, V.; Gidrol, X.; Balakirev, M. Y. High-Content Monitoring of Drug Effects in a 3D Spheroid Model. Front. Oncol. 2017, 7. 10.3389/fonc.2017.00293.

(13) Wang, Y.; Jeon, H. 3D Cell Cultures toward Quantitative High-Throughput Drug Screening. Trends Pharmacol. Sci. 2022, 43 (7), 569–581. 10.1016/j.tips.2022.03.014.

(14) Garnique, A. del M. B.; Parducci, N. S.; Miranda, L. B. L. de; Almeida, B. O. de; Sanches, L.; Machado-Neto, J. A.; Garnique, A. del M. B.; Parducci, N. S.; Miranda, L. B. L. de; Almeida, B. O. de; Sanches, L.; Machado-Neto, J. A. Two-Dimensional and Spheroid-Based Three-Dimensional Cell Culture Systems: Implications for Drug Discovery in Cancer. Drugs Drug Candidates 2024, 3 (2), 391–409. 10.3390/ddc3020024.

(15) Cha, Y. S.; Michaels, A.; Wang, J. Z.; Niu, Y.; Lin, Y.; Zhu, L.; Zhu, X.; Wang, K.; Murray, M.; Zhou, F. Recent Advances in 3D Cell Culture Models in Cancer Drug Development. J. Pharm. Investig. 2025, 55 (4), 557–573. 10.1007/s40005-025-00740-y.

(16) Trofimov, M. A.; Bulatov, I. P.; Lavrinenko, V. S.; Popov, V. E.; Petrova, V. S.; Bukatin, S.; Tyazhelnikov, S. F.; Trofimov, M. A.; Bulatov, I. P.; Lavrinenko, V. S.; Popov, V. E.; Petrova, V. S.; Bukatin, A. S.; Tyazhelnikov, S. F. Visualization, Data Extraction, and Multiparametric Analysis of 3D Pancreatic and Colorectal Cancer Cell Lines for High-Throughput Screening. Biomedicines 2026, 14 (1). 10.3390/biomedicines14010108.

(17) Bray, F.; Laversanne, M.; Sung, H.; Ferlay, J.; Siegel, R. L.; Soerjomataram, I.; Jemal, A. Global Cancer Statistics 2022: GLOBOCAN Estimates of Incidence and Mortality Worldwide for 36 Cancers in 185 Countries. CA. Cancer J. Clin. 2024, 74 (3), 229–263. 10.3322/caac.21834.

(18) Arutyunyan, I.; Jumaniyazova, E.; Makarov, A.; Fatkhudinov, T. In Vitro Models of Head and Neck Cancer: From Primitive to Most Advanced. J. Pers. Med. 2023, 13 (11), 1575. 10.3390/jpm13111575.

(19) Cancer: Principles & Practice of Oncology: Primer of the Molecular Biology of Cancer, 2nd Edition (Online Access Included). ProtoView 2015.

(20) Yilmaz, E.; Ismaila, N.; Bauman, J. E.; Dabney, R.; Gan, G.; Jordan, R.; Kaufman, M.; Kirtane, K.; McBride, S. M.; Old, M. O.; Rooper, L.; Saba, N. F.; Sheth, S.; Subramaniam, R. M.; Wise-Draper, T. M.; Wong, D.; Mell, L. K. Immunotherapy and Biomarker Testing in Recurrent and Metastatic Head and Neck Cancers: ASCO Guideline. J. Clin. Oncol. 2023, 41 (5), 1132–1146. 10.1200/JCO.22.02328.

(21) Moy, J. D.; Moskovitz, J. M.; Ferris, R. L. Biological Mechanisms of Immune Escape and Implications for Immunotherapy in Head and Neck Squamous Cell Carcinoma. Eur. J. Cancer Oxf. Engl. 1990 2017, 76, 152–166. 10.1016/j.ejca.2016.12.035.

(22) Bonomi, M.; Patsias, A.; Posner, M.; Sikora, A. The Role of Inflammation in Head and Neck Cancer. Adv. Exp. Med. Biol. 2014, 816, 107–127. 10.1007/978-3-0348-0837-8_5.

(23) Tong, C. C. L.; Kao, J.; Sikora, A. G. Recognizing and Reversing the Immunosuppressive Tumor Microenvironment of Head and Neck Cancer. Immunol. Res. 2012, 54 (1–3), 266–274. 10.1007/s12026-012-8306-6.

(24) Varilla, V.; Atienza, J.; Dasanu, C. A. Immune Alterations and Immunotherapy Prospects in Head and Neck Cancer. Expert Opin. Biol. Ther. 2013, 13 (9), 1241–1256. 10.1517/14712598.2013.810716.

(25) Liu, C.; Wang, M.; Zhang, H.; Li, C.; Zhang, T.; Liu, H.; Zhu, S.; Chen, J. Tumor Microenvironment and Immunotherapy of Oral Cancer. Eur. J. Med. Res. 2022, 27 (1), 198. 10.1186/s40001-022-00835-4.

(26) Harris, M.; Wang, X. G.; Jiang, Z.; Goldberg, G. L.; Casadevall, A.; Dadachova, E. Radioimmunotherapy of Experimental Head and Neck Squamous Cell Carcinoma (HNSCC) with E6-Specific Antibody Using a Novel HPV-16 Positive HNSCC Cell Line. Head Neck Oncol. 2011, 3 (1), 9. 10.1186/1758-3284-3-9.

(27) Utama, R. H.; Atapattu, L.; O’Mahony, A. P.; Fife, C. M.; Baek, J.; Allard, T.; O’Mahony, K. J.; Ribeiro, J. C. C.; Gaus, K.; Kavallaris, M.; Gooding, J. J. A 3D Bioprinter Specifically Designed for the High-Throughput Production of Matrix-Embedded Multicellular Spheroids. iScience 2020, 23 (10). 10.1016/j.isci.2020.101621.

(28) Utama, R. H.; Tan, V. T. G.; Tjandra, K. C.; Sexton, A.; Nguyen, D. H. T.; O’Mahony, P.; Du, E. Y.; Tian, P.; Ribeiro, J. C. C.; Kavallaris, M.; Gooding, J. J. A Covalently Crosslinked Ink for Multimaterials Drop-on-Demand 3D Bioprinting of 3D Cell Cultures. Macromol. Biosci. 2021, 21 (9), 2100125. 10.1002/mabi.202100125.

(29) Du, E. Y.; Jung, M.; Skhinas, J.; Tolentino, M. A. K.; Noy, J.; Jamshidi, N.; Houng, J. L.; Tjandra, K. C.; Engel, M.; Utama, R.; Tilley, R. D.; Kavallaris, M.; Gooding, J. J. 3D Bioprintable Hydrogel with Tunable Stiffness for Exploring Cells Encapsulated in Matrices of Differing Stiffnesses. ACS Appl. Bio Mater. 2023, 6 (11), 4603–4612. 10.1021/acsabm.3c00334.

(30) Chen, X.; O’Mahony, A. P.; Barber, T. J. Effect of 3D-Bioprinted Droplet Impact Dynamics on a Pre-Printed Soft Hydrogel Matrix. Exp. Fluids 2023, 64 (3), 60. 10.1007/s00348-023-03583-1.

(31) Yin, Y.; Vázquez-Rosado, E. J.; Wu, D.; Viswananthan, V.; Farach, A.; Farach-Carson, M. C.; Harrington, D. A. Microfluidic Coaxial 3D Bioprinting of Cell-Laden Microfibers and Microtubes for Salivary Gland Tissue Engineering. Biomater. Adv. 2023, 154, 213588. 10.1016/j.bioadv.2023.213588.

(32) Wadell, H. Volume, Shape, and Roundness of Rock Particles. J. Geol. 1932, 40 (5), 250–280.

(33) Chaves, P.; Garrido, M.; Oliver, J.; Pérez-Ruiz, E.; Barragan, I.; Rueda-Domínguez, A. Preclinical Models in Head and Neck Squamous Cell Carcinoma. Br. J. Cancer 2023, 128 (10), 1819–1827. 10.1038/s41416-023-02186-1.

(34) Chen, Y.; Wang, S.; Zheng, L.; Chen, L.; Xu, F.; Guo, S.; Meng, J. Generating Tumor-Specific T Cells Based on a Head and Neck Cancer Organoid for Adoptive Cell Therapy. Front. Immunol. 2025, 16, 1573965. 10.3389/fimmu.2025.1573965.

(35) Yoon, H.-N.; Kim, J.; Gu, D.; Lee, J.; Kim, S. Y.; Kim, H. J.; Jeong, J.; Shin, D.; Jung, Y.-S.; Chung, M. K.; Lee, S.-J.; Choi, S. Y. Patient-Derived 3D Organoid Platform for Functional Assessment of GPC3-Targeted CAR T Cell Cytotoxic Activity in Head and Neck Squamous Cell Carcinoma. Oral Oncol. 2026, 172, 107814. 10.1016/j.oraloncology.2025.107814.

(36) Graf, J.; Moore, D.; Grimes, C. L.; Fromen, C. A.; Kloxin, A. M. High-Throughput Bioprinted 3D Cultures for Probing Host–Pathogen Interactions in Bioinspired Microenvironments. RSC Appl. Polym. 2026. 10.1039/D5LP00285K.

(37) Jung, M.; Poltavets, V.; Skhinas, J. N.; Tax, G.; Kamili, A.; Xie, J.; Ghamrawi, S.; Graber, P.; Mao, J.; Wong-Erasmus, M.; Cui, L.; Kimpton, K.; Venkat, P.; Mayoh, C.; Lin, A.; Fleuren, E. D. G.; Fordham, A. M.; Barger, Z.; Grady, J.; Thomas, D. M.; Du, E. Y.; Graf, N. S.; Cowley, M. J.; Gifford, A. J.; Fletcher, J. I.; Lau, L. M. S.; Dolman, M. E. M.; Gooding, J. J.; Kavallaris, M. High-Throughput 3D Engineered Paediatric Tumour Models for Precision Medicine. Mol. Syst. Biol. 2025, 21 (12), 1748–1777. 10.1038/s44320-025-00152-y.

(38) Decarli, M. C.; Seijas-Gamardo, A.; Rademakers, T.; Sanchez, A. A.; Wieringa, P.; Vicente L Silva, J.; Moraes, Â. M.; Moroni, L.; Mota, C. An Automated Pipeline for Tracking and Measuring Cell Spheroids Encapsulated in 3D Hydrogel Systems. Biofabrication 2026, 18 (2), 025009. 10.1088/1758-5090/ae4893.

(39) Mohammadrezaei, D.; Moghimi, N.; Vandvajdi, S.; Powathil, G.; Hamis, S.; Kohandel, M. Predicting and Elucidating the Post-Printing Behavior of 3D Printed Cancer Cells in Hydrogel Structures by Integrating in-Vitro and in-Silico Experiments. Sci. Rep. 2023, 13 (1), 1211. 10.1038/s41598-023-28286-9.

(40) Darling, N. J.; Hung, Y.-S.; Sharma, S.; Segura, T. Controlling the Kinetics of Thiol-Maleimide Michael-Type Addition Gelation Kinetics for the Generation of Homogenous Poly(Ethylene Glycol) Hydrogels. Biomaterials 2016, 101, 199–206. 10.1016/j.biomaterials.2016.05.053.

(41) Jansen, L. E.; Negrón-Piñeiro, L. J.; Galarza, S.; Peyton, S. R. Control of Thiol-Maleimide Reaction Kinetics in PEG Hydrogel Networks. Acta Biomater. 2018, 70, 120–128. 10.1016/j.actbio.2018.01.043.

(42) Richbourg, N. R.; Rampal, A.; Lorenzana, A.; Flechas-Beltran, J. F.; Peyton, S. R. The Intersecting Physical Mechanisms That Regulate Cell Viability in 3D Synthetic Hydrogels. Adv. Mater. n/a (n/a), e21245. 10.1002/adma.202521245.

(43) Micalet, A.; Moeendarbary, E.; Cheema, U. 3D In Vitro Models for Investigating the Role of Stiffness in Cancer Invasion. ACS Biomater. Sci. Eng. 2023, 9 (7), 3729–3741. 10.1021/acsbiomaterials.0c01530.

(44) Graf, J.; Moore, D.; Grimes, C. L.; Fromen, C. A.; Kloxin, A. M. High-Throughput Bioprinted 3D Cultures for Probing Host–Pathogen Interactions in Bioinspired Microenvironments. RSC Appl. Polym. 2026. 10.1039/D5LP00285K.

(45) Dean, T.; Li, N. T.; Cadavid, J. L.; Ailles, L.; McGuigan, A. P. A TRACER Culture Invasion Assay to Probe the Impact of Cancer Associated Fibroblasts on Head and Neck Squamous Cell Carcinoma Cell Invasiveness. Biomater. Sci. 2020, 8 (11), 3078–3094. 10.1039/C9BM02017A.

