## SupplementalInformation for "3D Droplet-Based Bioprinting of Customized *In Vitro* Head and Neck Cancer Tumor Microenvironment Models"

**Supporting Information:**

Immunofluorescence (IF) staining detailed methods:

Samples at day 7 of culture were prepared for IF staining by aspirating cell culture media from each well, and incubating with DPBS for 5 min / 25 °C. All DPBS rinse steps were conducted under light orbital shaking. DPBS was aspirated from each well, 100 μL 4% (w/v) paraformaldehyde (PFA) (97%, Alfa Aesar A11313) in DPBS was added to each well and incubated for 20 min / 25°C. Fixative was aspirated, each hydrogel was incubated with DPBS 3x5 min/ 25°C, then permeabilized with permeabilized with 0.1% (v/v) Triton X-100 (Thermo Scientific A16046.AP) for 20 min / 25°C. The permeabilization buffer was aspirated, and each hydrogel was incubated with DPBS 3x5 min/ 25°C . Samples were incubated with 100 μL of blocking buffer (10% (v/v) normal goat serum (Millipore, S26-100ML) in PBS) overnight at 4°C. Blocking buffer was aspirated, and 100 μL of primary antibody solutions, diluted in blocking buffer to concentrations noted in Table SI-1, were added to each well. Multiwell plates were covered in aluminum foil and incubated at 4˚C overnight for 16-22 hours with light orbital shaking at 60 rpm. Primary antibody solutions were aspirated, and wells were washed 3x5 min with 0.1% (v/v) Tween® 20 (Fisher BioReagents BP337-500) in DPBS. Secondary antibodies were diluted to concentrations noted in Table SI-1 using 0.1% (v/v) Triton X-100 in DPBS including 1 μg/μL DAPI (4', 6-diamidino-2-phenylindole, thermoscientific #62247). 100 μL of secondary antibody solutions were added to each well, and multiwell plates were covered in aluminum foil and incubated at 4˚C overnight for 16-22 hours with light orbital shaking at 60 rpm. Secondary antibodies were aspirated, and wells were incubated with DPBS 3x5 min/ 25°C. Final stained samples were stored in 100 μL 0.1% sodium azide (Acros Organics B0143741) in DPBS and imaged.

Table SI-1: Immunostaining parameters

| Target | Host | Isotype | Company | Catalog # | Lot # | Stock Conc. (mg/mL) | Working Dilution (v/v) |
|---|---|---|---|---|---|---|---|
| **Primary Antibodies** | | | | | | | |
| fibronectin | Rb | polyclonal | abcam | AB2413 | 1111584-1 | 0.024 | 1:200 |
| vimentin | Rb | IgG | abcam | AB92547 | 1072376-91 | 0.261 | 1:400 |
| Ki67 | Rb | IgG | abcam | AB16667 | 1090780-150 | 0.031 | 1:250 |
| pan-cytokeratin | Ms | IgG1 | Invitrogen | 41-9003-82 | 3175289 | 0.2 | 1:100 |
| EGFR | Ms | IgG1 | Invitrogen | MA5-13070 | 79660881 | 0.2 | 1:200 |
| laminin | Rb | polyclonal | Novus Biologicals | NB300-144 | 4267502 | 1 | 1:200 |
| E-cadherin | Rb | IgG | Cell Signaling Technology | 3195T | 15 | 0.054 | 1:200 |
| N-cadherin | Ms | IgG1 | BD Transduction Laboratories | 610920 | 3324728 | 0.25 | 1:200 |
| EpCAM | Ms | IgG1 | Cell Signaling Technology | 2929S | 11 | N/A | 1:200 |
| **Secondary Antibodies** | | | | | | | |
| Goat anti-mouse IgG (H+L) Highly Cross-Adsorbed Secondary Antibody Alexa Fluor™ 488 | Gt | polyclonal | Invitrogen | A-11029 | | 2 | 1:500 |
| Goat anti-rabbit IgG (H+L) Cross-Adsorbed Secondary Antibody Alexa Fluor™ 568 | Gt | polyclonal | Invitrogen | A-11011 | | 2 | 1:1000 |

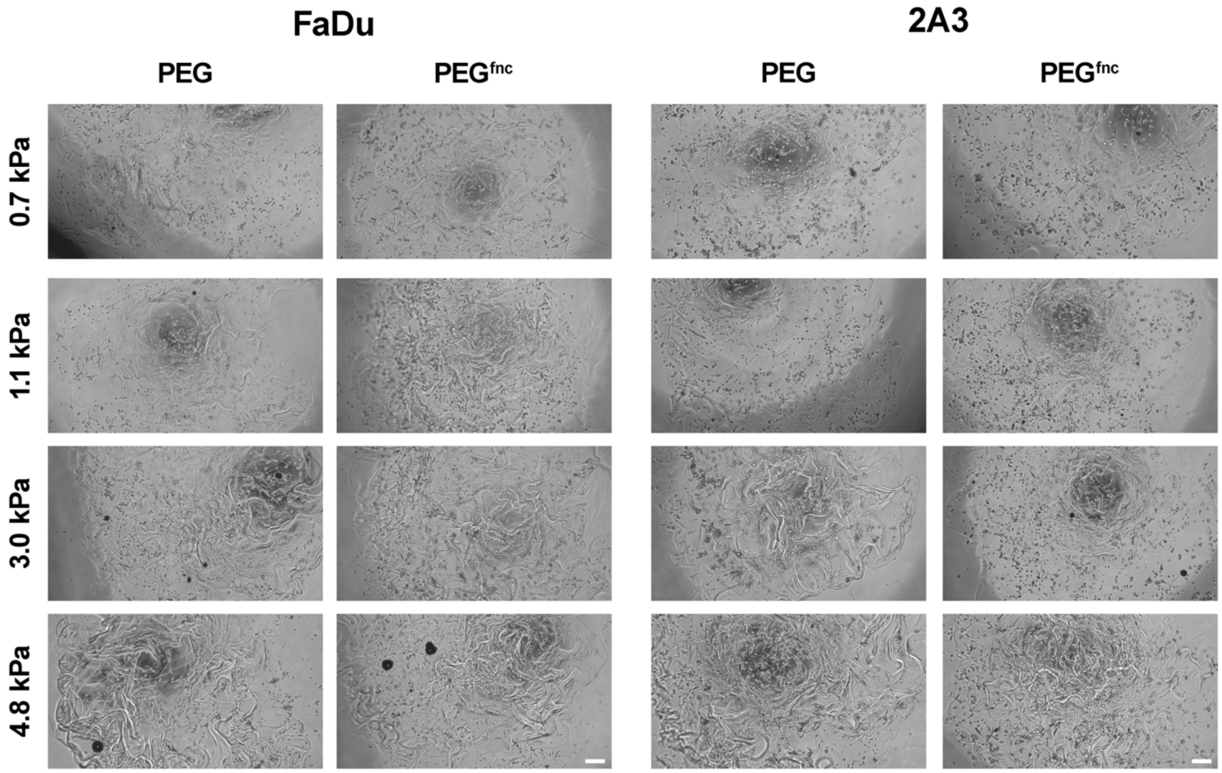


**Figure SI-1.** Light microscopy images shown above were recorded within the first 24 hours after printing (day 0) for each cell type in each hydrogel condition. Cells were thoroughly resuspended in hydrogel precursor solutions prior to printing. Images demonstrate single-cell conditions in nearly every case, with minimal association at this early timepoint. Scale bar = 100 µm.

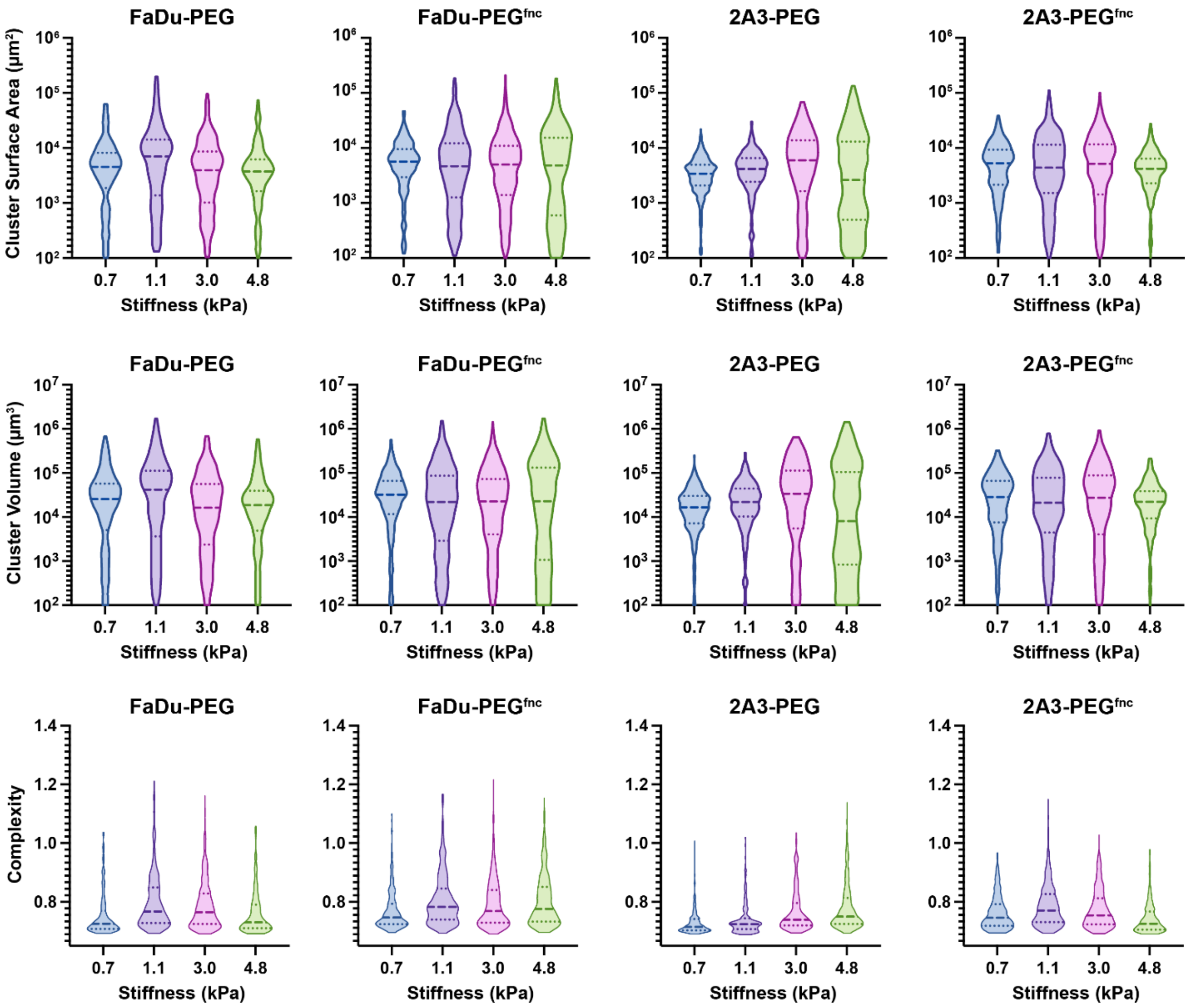


**Figure SI-2.** Violin plots for FaDu and 2A3 cluster measurements of surface area, volume, and complexity. Each violin represents measurements from the total of all clusters across triplicate samples, ranging from n=237-670. Dashed lines represent the median value, and dotted lines represent the upper and lower quartile boundary. Violin boundaries for area and volume were calculated from log-transform values and plotted on a linear antilog scale. Complexity was calculated from area and volume measures, as noted in Methods, and plotted on a linear scale. Complexity of 0.684 represents a perfect sphere, and any values above that reflect further deviations from a perfect sphere.

**Table SI-2**: Term Assessment for Model Fitting

| | | p-values for significance | | |
|---|---|---|---|---|
| | **Tested Interaction Terms** | **Area** | **Volume** | **Complexity** |
| 4-way | log_stiffness:celltype:matrix:time | 0.1322 | 0.0945 | 3.651E-6 |
| 3-way | log_stiffness:celltype:matrix | 0.1164 | 0.3273 | n/a |
| | log_stiffness:celltype:time | 0.9569 | 0.9248 | n/a |
| | log_stiffness:matrix:time | 0.2577 | 0.4082 | n/a |
| | celltype:matrix:time | 0.2798 | 0.272 | n/a |
| 2-way | log_stiffness:celltype | 0.0134 | 0.02875 | n/a |
| | log_stiffness:matrix | 0.3017 | 0.1326 | n/a |
| | log_stiffness:time | 0.007 | 0.00323 | n/a |
| | celltype:matrix | 0.0851 | 0.9748 | n/a |
| | celltype:time | 8.6E-8 | 2.41E-5 | n/a |
| | matrix:time | 0.0145 | 0.04176 | n/a |
